# Maximal coral thermal tolerance is found at intermediate diel temperature variability

**DOI:** 10.1101/2023.03.27.534434

**Authors:** Kristen T. Brown, Marcelina Martynek, Katie L. Barott

## Abstract

1. It has become critically important to identify environmental drivers of enhanced thermal tolerance in coral populations as ocean warming threatens the persistence of coral reef ecosystems globally. Variable temperature regimes that expose corals to sub-lethal heat stress have been recognized as a mechanism to increase coral thermotolerance and lessen coral bleaching; however, there is a need to better understand which thermal regimes are best for promoting coral stress hardening, and if thermal priming results in consistent benefits across species with distinct life-history strategies.
2. Standardized thermal stress assays were used to determine the relative thermal tolerance of three divergent genera of corals (*Acropora, Pocillopora* and *Porites*) originating from six reef sites fluctuating in temperature by up to 7.7°C day^-1^, with an annual mean diel variability of 1–3°C day^-1^. Bleaching severity and dark-acclimated photochemical yield (F_v_/F_m_) were quantified following exposure to five temperature treatments ranging from 23.0 to 36.3°C — up to 9°C above the regional maximum monthly mean.
3. The greatest thermal tolerance across all species was found at the site with intermediate mean diel temperature variability (2.2°C day^-1^), suggesting there is an optimal priming exposure that leads to maximal thermotolerance. Interestingly, *Acropora* and *Pocillopora* originating from the least thermally variable regimes (i.e., <1.3°C day^-1^) had lower thermal tolerance than corals from the most variable sites (i.e., > 2.8°C day^-1^), whereas the opposite was true for *Porites*, suggesting divergent responses to priming across taxa.
4. We highlight that fine-scale heterogeneity in temperature dynamics across habitats can increase coral thermal tolerance in diverse coral lineages, although in a non-linear manner. Remarkably, comparisons across global studies revealed that the range in coral thermotolerance uncovered in this study across a single reef system (<5 km) were as large as differences observed across vast latitudinal gradients (>300 km). This important finding indicates that local gene flow could improve thermal tolerance between habitats. However, as climate change continues, exposure to intensifying marine heatwaves is already compromising thermal priming as a mechanism to enhance coral thermal tolerance and bleaching resistance.

## Introduction

Ocean warming due to anthropogenic greenhouse gas emissions is the greatest threat to the persistence of coral reefs in the Anthropocene (Hoegh-Guldberg et al., 2019). Reef-building corals live at the upper edge of their thermal limits, and persistent temperatures just 1°C above a coral’s typical summer maximum can cause the breakdown of the symbiosis between the coral and its endosymbiotic algae (family Symbiodiniaceae) — a phenomenon known as coral bleaching (van Woesik et al., 2022). Marine heatwaves resulting in mass coral bleaching are now occurring globally on multi-decadal time-scales, having gone from up to one mild event a decade last century to as many as five per decade in modern times (Hughes et al., 2018). It has thus become increasingly important to identify thermally tolerant coral populations capable of surviving intensifying marine heatwaves. Encouragingly, coral populations with elevated heat tolerance have been found within various thermally extreme environments, such as mangrove lagoons and tidally-dominated reef flats, which expose inhabitants to short-term temperature extremes not predicted to occur on ordinary reefs until 2100 (Brown et al., 2022; Camp et al., 2019; Schoepf et al., 2015). Life-long exposure to high diel temperature variability has thus emerged as an important factor in promoting elevated coral bleaching thresholds (Barshis et al., 2013; Kenkel & Matz, 2016; Oliver & Palumbi, 2011a; Palumbi et al., 2014; Voolstra et al., 2020). However, we still lack a clear understanding of the priming exposure (e.g., magnitude of thermal variability) most beneficial for coral stress hardening (Hackerott et al., 2021), as not all variable temperature regimes maximize coral thermotolerance (Klepac & Barshis, 2022; Schoepf et al., 2019).

A comprehensive understanding of how environmental drivers influence coral thermal tolerance requires a direct comparison of coral genera with different life-history strategies (e.g., competitive v. stress-tolerant) (Darling et al., 2012) and evolutionary histories (Kitahara et al., 2010). For example, the same highly variable habitats that promote elevated thermotolerance in competitive *Acropora hyacinthus* do not lead to elevated thermotolerance in stress-tolerant *Porites lobata* (Klepac & Barshis, 2020; Palumbi et al., 2014; Thomas et al., 2018). Similarly, across larger spatial scales, relative thermotolerance is not predictable across corals with distinct life-history strategies. For example, competitive corals (*Acropora hemprichii*) were identified as the most thermally tolerant when compared to weedy (*Pocillopora verrucosa*; *Stylophora pistillata*) or stress-tolerant (*Porites lobata*) taxa across the Red Sea, whereas across the Coral Sea (eastern Australia), competitive acroporids were the least thermally tolerant when compared to weedy species (*P. verrucosa; P. meandrina*) (Marzonie et al., 2022).

Discrepancies may be due to experimental design (Grottoli et al., 2021), fine-scale differences in Symbiodiniaceae genotypes (Oliver & Palumbi, 2011b), seasonality (Berkelmans & Willis, 1999), and/or increasing history of heat stress that may compromise thermal priming as a protective mechanism (Ainsworth et al., 2016; Klepac & Barshis, 2020; Schoepf et al., 2015). To accurately evaluate thermal tolerance and predict the impact of climate change on coral reef ecosystems, standardized comparisons across diverse species and environmental mosaics are critically needed (Grottoli et al., 2021; Voolstra et al., 2020).

Here, we employed a standardized experimental heat stress assay (e.g., (Voolstra et al., 2020)) to determine the relative thermotolerance of three coral genera across Heron Island, southern Great Barrier Reef (GBR). Corals were collected from six sites encompassing five distinct geomorphological habitats (Phinn et al., 2012), which differ in long-term thermal variability, fluctuating by up to 7.7°C day^-1^ (Brown et al., 2022). These ranges are comparable to other study systems across the globe, including: the highly variable pools of Ofu, American Sāmoa (up to 6°C day^-1^) (Thomas et al., 2018), exposed and protected sites of the central Red Sea (up to 6.5°C day^-1^) (Voolstra et al., 2020), intertidal and subtidal environments of the Kimberley in western Australia (up to 7°C day^-1^) (Schoepf et al., 2015), and mangrove lagoons of the GBR (7.7°C day^-1^) (Camp et al., 2019). Corals representing three distinct life-history strategies and the two clades of Scleractinia were investigated — *Acropora* cf. *aspera* (competitive; Complexa), *Pocillopora* cf. *damicornis* (weedy; Robusta) and *Porites* cf. *lobata* (stress-tolerant; Complexa) — to explore if thermal priming results in consistent benefits across species.

Thermal tolerance was compared to coral community resilience in the aftermath of a recent marine heatwave (Brown et al., 2022) to see if increased thermotolerance is protective during modern marine heatwaves. Finally, this standardized experimental approach allowed us to compare coral thermal thresholds across studies from different reef systems and across large spatial scales.

## Materials and Methods

### Study location

This study was conducted across six sites at Heron Island, southern Great Barrier Reef (23°27’ S 151°55’ E; Figure 1), which included at least one representative from each geomorphological habitat of Heron Reef (site; depth): reef slope [Fourth Point (FP; 4.2 m); Harry’s Bommie (HB; 6.1 m)], reef crest (RC; 0.9 m), reef flat (RF; 0.7 m), shallow lagoon (SL; 1.3 m) and deep lagoon (DL; 2.6 m) (Phinn et al., 2012) (Figure 1). Historically, hard coral cover has been greatest within the reef slope (∼60%), which is dominated by corals of the family Acroporidae, whereas within the lagoon, sites have peaked around 20% coral cover and are principally composed of *Pocillopora* and massive *Porites* (Brown et al., 2022; Connell et al., 1997; Roelfsema et al., 2021). Experiments were performed in the austral spring to avoid any potentially confounding thermal stress that is becoming increasingly common during the summer (Marzonie et al., 2022). While absolute thermal tolerance can differ across seasons (Berkelmans & Willis, 1999), relative thermal tolerance between individuals across seasons remains consistent (Cunning et al., 2021; Evensen et al., 2022)

**Figure 1.**
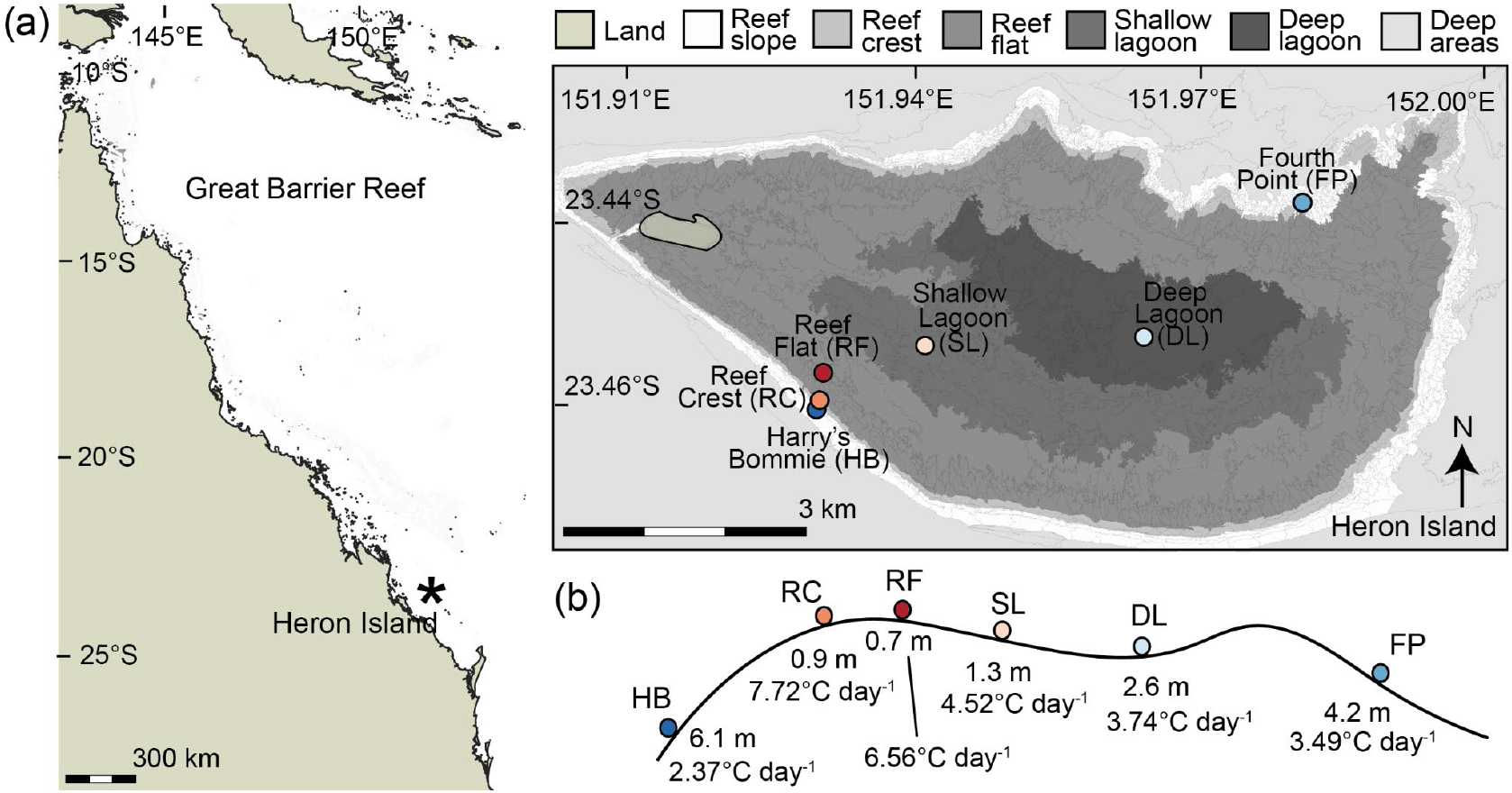
Study location and reef profile of Heron Island, southern Great Barrier Reef. (a) Map of the Great Barrier Reef, with an asterisk showing the location of Heron Island. Map inset details the geomorphological habitats of Heron Reef, redrawn from the data of (Phinn et al., 2012), with sampling sites indicated. (b) Reef profile showing the depth and maximum diel temperature variability across the reef sites investigated in this study.

### Evaluation of temperature variability

Seawater temperatures were recorded hourly from July 2015 to September 2022 (Figure 2). From July 2015 – November 2016, seawater temperatures were recorded by use of Conductivity Temperature Depth units (CTD; SBE 16plus V2 SEACAT) until their removal, at which time cross-calibrated HOBO Pendant loggers (UA-001-64; ± 0.552°C accuracy) were deployed. Logger accuracy was assessed at the end of each deployment period using a water bath (Thermo Scientific Precision TSGP20). Temperature dynamics (e.g., mean, maximum, diel variability) were calculated at each site across the one-year period from September 2015 – August 2016, as this period included the most complete record across all sites and did not include a marine heatwave (Table 1).

**Figure 2.**
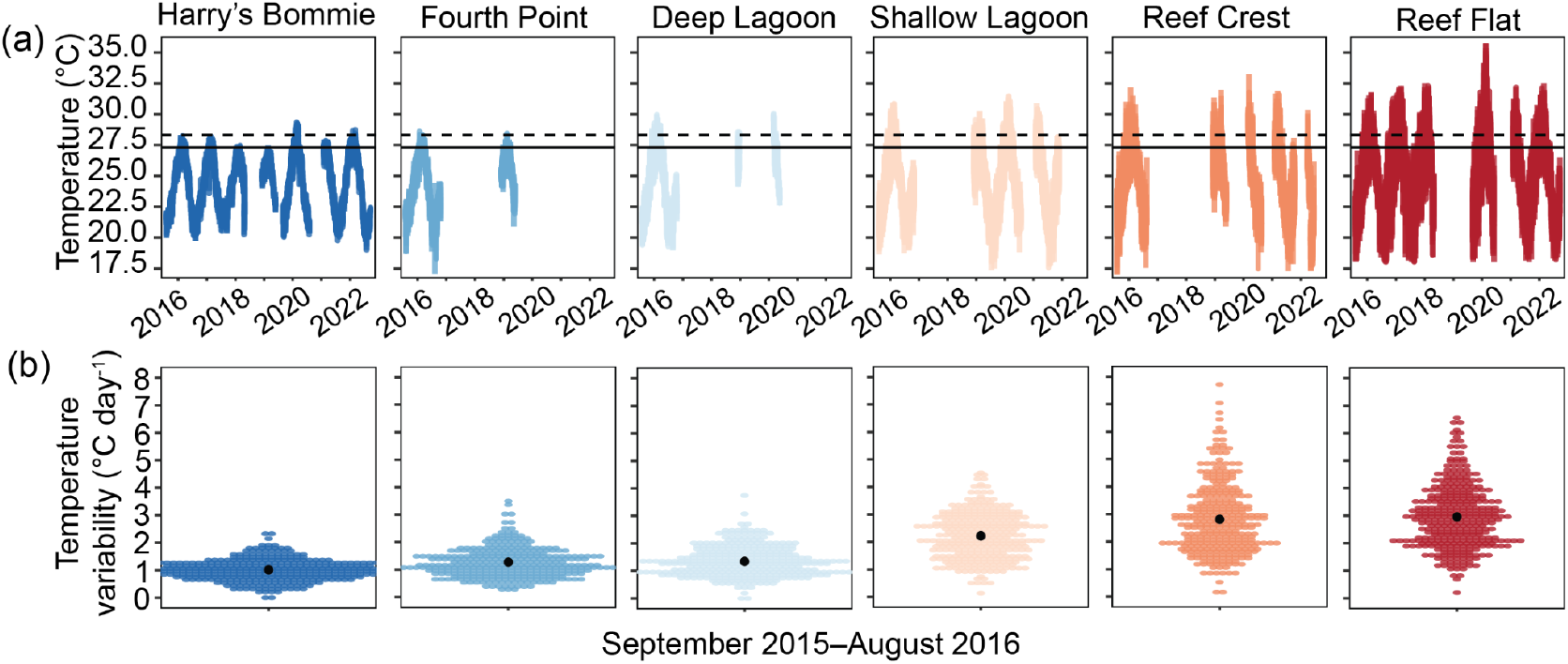
Temperature profiles across Heron Reef from least to most thermally variable. (a) Hourly seawater temperatures were recorded from August 2015 to September 2022. Solid horizontal line indicates the region’s climatological maximum monthly mean (MMM; 27.3°C) and dashed horizontal line indicates the region’s coral bleaching threshold (MMM + 1°C; 28.3°C). (b) Diel temperature variability across September 2015–August 2016, where individual points represent each day and the black point indicates the mean across the year.

**Table 1.**
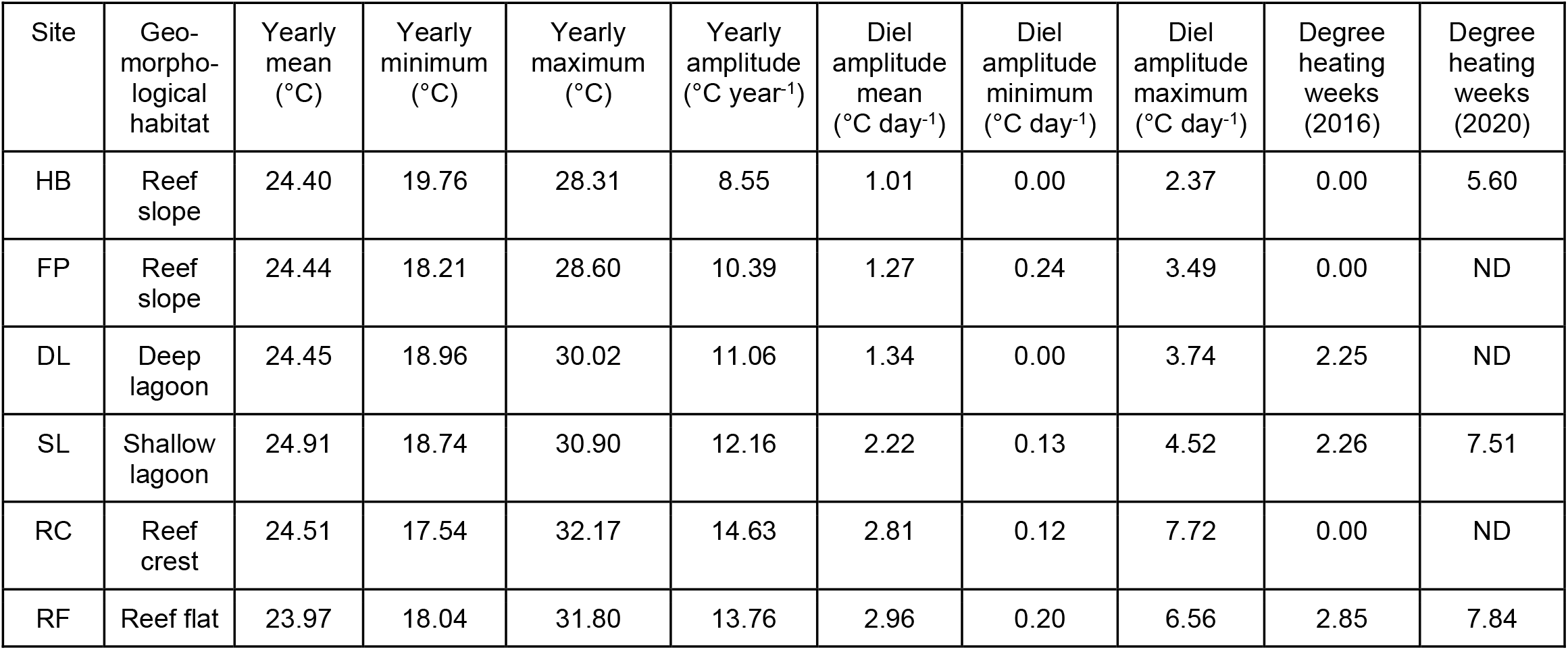
Temperature metrics across Heron Island, southern Great Barrier Reef for the year-long period of September 2015 – August 2016. Sites are listed in order from least to most thermally variable and are indicated by abbreviation, where: HB = Harry’s Bommie, FP = Fourth Point, DL = Deep Lagoon, SL= Shallow Lagoon, RC = Reef Crest, and RF = Reef Flat. ND indicates no data.

| Site | Geo-<br>morpho-<br>logical<br>habitat | Yearly<br>mean<br>(°C) | Yearly<br>minimum<br>(°C) | Yearly<br>maximum<br>(°C) | Yearly<br>amplitude<br>(°C year<sup>-1</sup>) | Diel<br>amplitude<br>mean<br>(°C day<sup>-1</sup>) | Diel<br>amplitude<br>minimum<br>(°C day<sup>-1</sup>) | Diel<br>amplitude<br>maximum<br>(°C day<sup>-1</sup>) | Degree<br>heating<br>weeks<br>(2016) | Degree<br>heating<br>weeks<br>(2020) |
| --- | --- | --- | --- | --- | --- | --- | --- | --- | --- | --- |
| HB | Reef<br>slope | 24.40 | 19.76 | 28.31 | 8.55 | 1.01 | 0.00 | 2.37 | 0.00 | 5.60 |
| FP | Reef<br>slope | 24.44 | 18.21 | 28.60 | 10.39 | 1.27 | 0.24 | 3.49 | 0.00 | ND |
| DL | Deep<br>lagoon | 24.45 | 18.96 | 30.02 | 11.06 | 1.34 | 0.00 | 3.74 | 2.25 | ND |
| SL | Shallow<br>lagoon | 24.91 | 18.74 | 30.90 | 12.16 | 2.22 | 0.13 | 4.52 | 2.26 | 7.51 |
| RC | Reef<br>crest | 24.51 | 17.54 | 32.17 | 14.63 | 2.81 | 0.12 | 7.72 | 0.00 | ND |
| RF | Reef flat | 23.97 | 18.04 | 31.80 | 13.76 | 2.96 | 0.20 | 6.56 | 2.85 | 7.84 |

**Table 2.** Replication statement with the key elements of experimental design and replication.

| Scale of inference | Scale at which the factor of interest is applied | Number of replicates at the appropriate scale |
| --- | --- | --- |
| Reef | Sites | 6 sites encompassing each geomorphological habitat of Heron Reef |
| Species | Species | 3 species: 1 competitive species (*Acropora* cf. *aspera*), 1 weedy species (*Pocillopora* cf. *damicornis*), 1 stress-tolerant species (*Porites* cf. *lobata*) |
| Individuals | Treatment | 5–10 individuals for each site × species × treatment combination |

### Sample collection

Three morphologically distinct coral species with distinct life-history strategies were examined: *Acropora* cf. *aspera* (competitive; branching open), *Pocillopora* cf. *damicornis* (weedy; branching closed), and *Porites* cf. *lobata* (stress-tolerant; massive) (Darling et al., 2012) (Figure 3). Coral fragments were collected from 27 September to 6 October 2022 (Table S1). Ten colonies of each genus were sampled from each site except where noted (n = 5–10 colonies per species per site; Figure 3). *A*. cf. *aspera* was not collected at the Shallow Lagoon and Deep Lagoon as it was absent or rare. Following collection, corals were transported to Heron Island Research Station (HIRS) and placed in outdoor, flow-through seawater troughs under ambient temperatures (22.86 ± 0.02°C) until experimentation. Each colony was divided into five fragments (i.e., genetic clones) of ∼5 cm using bone cutters (*Acropora* and *Pocillopora*) or a brick saw (*Porites*). *Acropora* and *Pocillopora* were then suspended within the experimental tanks using fishing line. *Porites* fragments were placed on plastic grating at the bottom of the experimental tanks. Coral thermal tolerance experiments were initiated within 3–48 hours of collection (Table S1).

**Figure 3.**
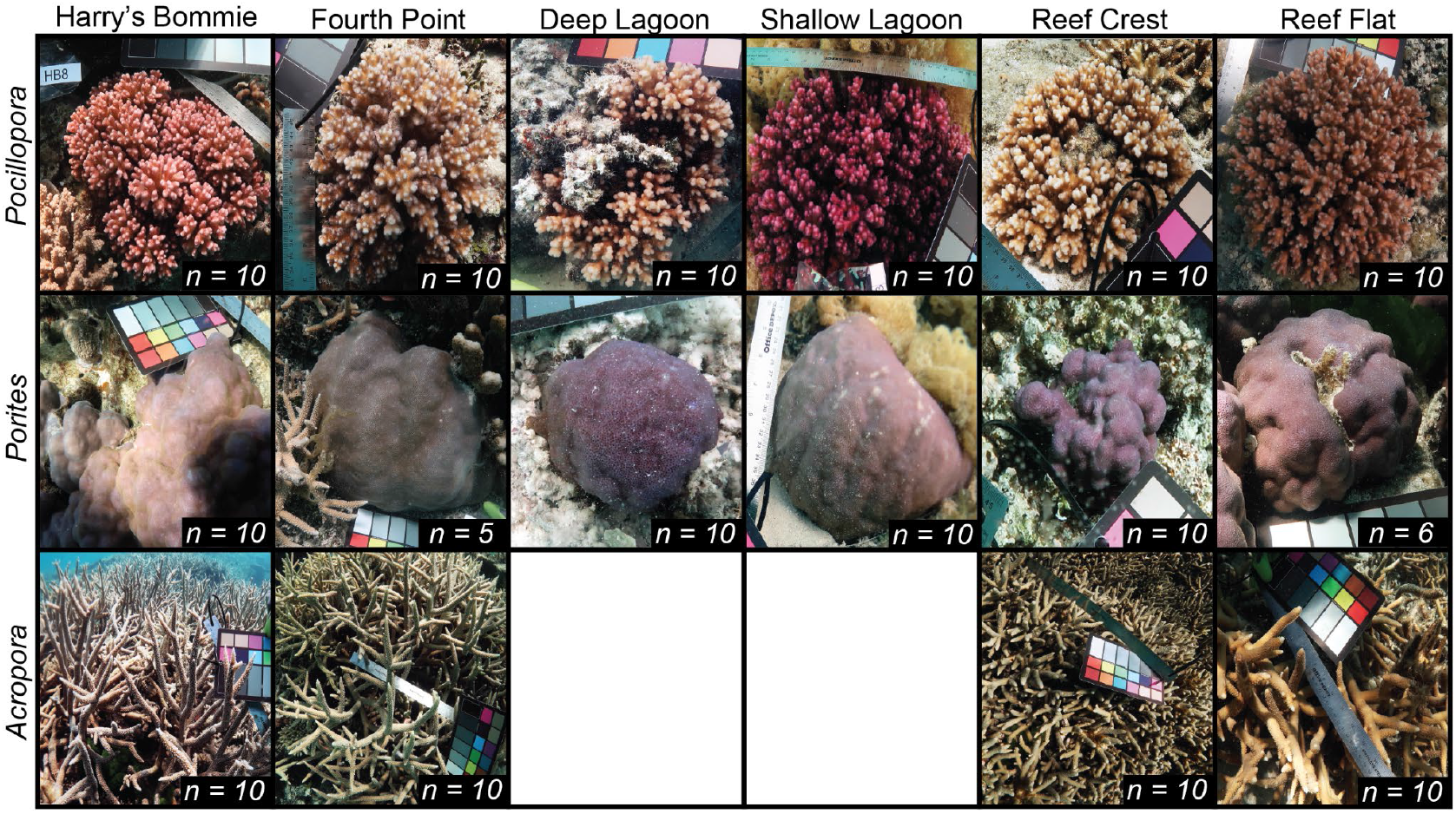
Representative images of coral genera across sites at Heron Island, southern Great Barrier Reef. Inset indicates the number of colonies collected at each site. At the Deep Lagoon and Shallow Lagoon, *Acropora* was absent or rare so was not evaluated.

### Acute heat stress experiment

A standardized temperature profile was used to measure heat tolerance in corals (e.g., (Cunning et al., 2021; Marzonie et al., 2022; Voolstra et al., 2020)) with minor modification. The climatological maximum monthly mean (MMM) of Heron Reef is 27.3°C (Weeks et al., 2008). A pilot experiment with all three genera indicated no difference in F_v_/F_m_ between MMM, MMM+3°C and MMM+6°C, so the latter two treatments were increased such that the five treatments used here included ambient, MMM, MMM+4°C, MMM+6.5°C, and MMM+9°C. Generally, experiments began at ∼12:00 with a 3-hr ramp to respective treatment temperatures (23°C, 27.3°C, 31.3°C, 33.8°C, 36.3°C), a 3-hr hold, and a 1-hr ramp down to MMM (Cunning et al., 2021; Marzonie et al., 2022; Voolstra et al., 2020) (Figure S1). Lights were turned off at the onset of the 1-hr ramp down to correspond with sunset. Due to experimental constraints (space, equipment, and time), only two treatments (n = 1 tank) were performed per day and each site was done in isolation.

Accordingly, a complete assay took two days per site, with treatments tested each day selected randomly (Table S1, Figure S1). A fragment from each coral colony was randomly placed into each treatment, so that all genotypes were present in each treatment. Temperatures were controlled using an Apex (Neptune Systems). Apex temperature probes were calibrated against a high-precision temperature probe (accuracy: ±0.4°C; HANNA HI-98190) at the onset of the experiment. Temperatures were also recorded using cross-calibrated temperature loggers (accuracy: ± 0.29°C; HOBO UA-001-64, Onset Computer Corporation). Photosynthetically active radiation (PAR) was static and controlled using aquarium lights (NICREW HyperReef LED, Shenzhen NiCai Technology Co.), averaging 250 μmol m^-2^ sec^-1^.

### Physiological responses to acute heat stress

At the end of the ramp and after 1 h of darkness (∼19:00), corals were assessed for dark-adapted photochemical yield (F_v_/F_m_) using a Diving-PAM (Walz GmbH) 5-mm diameter fiber-optic probe at a standardized distance (5 mm) above the coral tissue after F_0_ stabilized. Two random spots were measured on each fragment to obtain average measures of F_v_/F_m_. All readings with F_0_ values that were less than 110 were removed to avoid any false detections (Marzonie et al., 2022). The following morning at 07:00 corals were photographed with a color standard (WDKK Waterproof Color Chart, DGK Color Tools) to assess the effect of temperature on coral color, a proxy for relative chlorophyll density and bleaching severity (Voolstra et al., 2020; Winters et al., 2009) (for full details see Supplementary Methods).

### Statistical analyses

All statistical analyses were conducted using R software version 4.0.3 (R Core Team, 2021), and graphical representations were produced using *ggplot2* (Wickham, 2016). Differences in seawater temperature metrics were explored between sites (six levels: HB, FP, DL, SL, RC, RF) using linear models. Similarly, differences in temperature profiles (five levels: ambient, MMM, MMM+4°C, MMM+6.5°C, MMM+9°C) and experimental assays (n= 6 per temperature; see Table S1) were explored using a linear model. To assess for differences in coral color and photochemical yield between sites, treatments, and genera (three levels: *Acropora, Pocillopora, Porites*), linear mixed effects (lme) models were used, with genotype as a random effect. For all models, the Anova function in the package *car* was used to determine the significance of fixed effects and their interactions, with type II error structures applied for models that were not suggestive of interactions, and type III for models that were (Fox et al., 2012). Significant interactive effects were followed by pairwise comparison of estimate marginal means using the *emmeans* package with Tukey HSD adjusted *p* values (Lenth et al., 2018). Data were tested for homogeneity of variance and normality of distribution through graphical analyses of residual plots for all models.

To determine how heat tolerance differed amongst genera and sites, three-parameter log-logistic dose-response curves were fit to the median photochemical yield (F_v_/F_m_) across temperature treatments using the function drm in the package *drc* (Ritz et al., 2015). From these curves, the effective temperature to induce a 50% loss in F_v_/F_m_ (effective dose 50; ED50) was obtained following the methodology of (Evensen et al., 2022). Further, generalized additive models (GAMs) were fit to the median color score. Finally, to explore the relative influence of seawater temperature metrics (e.g., mean, maximum, mean daily amplitude; see Table 1 for all metrics) on coral thermal tolerance, GAMs were fit to allow for any possible non-linear effects (for full details see Supplementary Methods).

## Results

### Temperature variability differed across reef sites

Multiple years (2015–2022) of *in situ* temperature data demonstrated that daily (24-hr) mean (F= 4.1, p= 0.001), diel temperature amplitude (F= 1446.7, p< 0.0001), maximum temperatures (F= 91.9, p< 0.0001), and minimum temperatures (F= 40.1, p< 0.0001) significantly differed amongst sites (Figure 2). However, because an identical record was not obtained across all sites during the seven-year period, these significant differences were potentially driven by the patchiness of the time series. Therefore, the one-year period for which the most complete temperature record was obtained (September 2015–August 2016) was used for a more rigorous comparison of temperature dynamics across the six sites. During this period, daily mean temperatures were not significantly different across sites (F= 0.35, p= 0.88) (Table 1). In contrast, sites significantly differed in thermal variability (F= 338.1, p< 0.0001) (Figure 2), in order from the least to most variable (mean°C day^-1^ ± SE): Harry’s Bommie (1.01 ± 0.05), Fourth Point (1.27 ± 0.05), Deep Lagoon (1.34 ± 0.05), Shallow Lagoon (2.22 ± 0.05), Reef Crest (2.81 ± 0.05), and Reef Flat (2.96 ± 0.05) (Figure 2, Table 1). Pairwise comparisons revealed significant differences in mean daily temperature amplitude between all sites (p < 0.0006), except the Reef Crest and Reef Flat (p = 0.22) and Deep Lagoon and Fourth Point (p = 0.89) (Figure 2). Maximum diel temperature variability followed similar patterns, in order from the least to most variable (°C day^-1^): Harry’s Bommie (2.37), Fourth Point (3.49), Deep Lagoon (3.74), Shallow Lagoon (4.52), Reef Flat (6.56) and Reef Crest (7.72) (Figure 1, Table 1). Extreme temperature incursions leading to variability above 5°C day^-1^ were only observed at Reef Crest and Reef Flat, with the most extreme ranges observed at the Reef Crest (up to 7.7°C day^-1^), yet the highest frequency of extreme values occurred at the Reef Flat (Table 1, Figure 2).

### Relative thermal tolerance under acute heat stress

Target temperature profiles were successfully attained across the experimental heat stress assays (Figure S1). Importantly, there were no significant differences in temperature profiles of corresponding treatments between assays (F= 0.41, p= 0.84) (Figure 4a).

**Figure 4.**
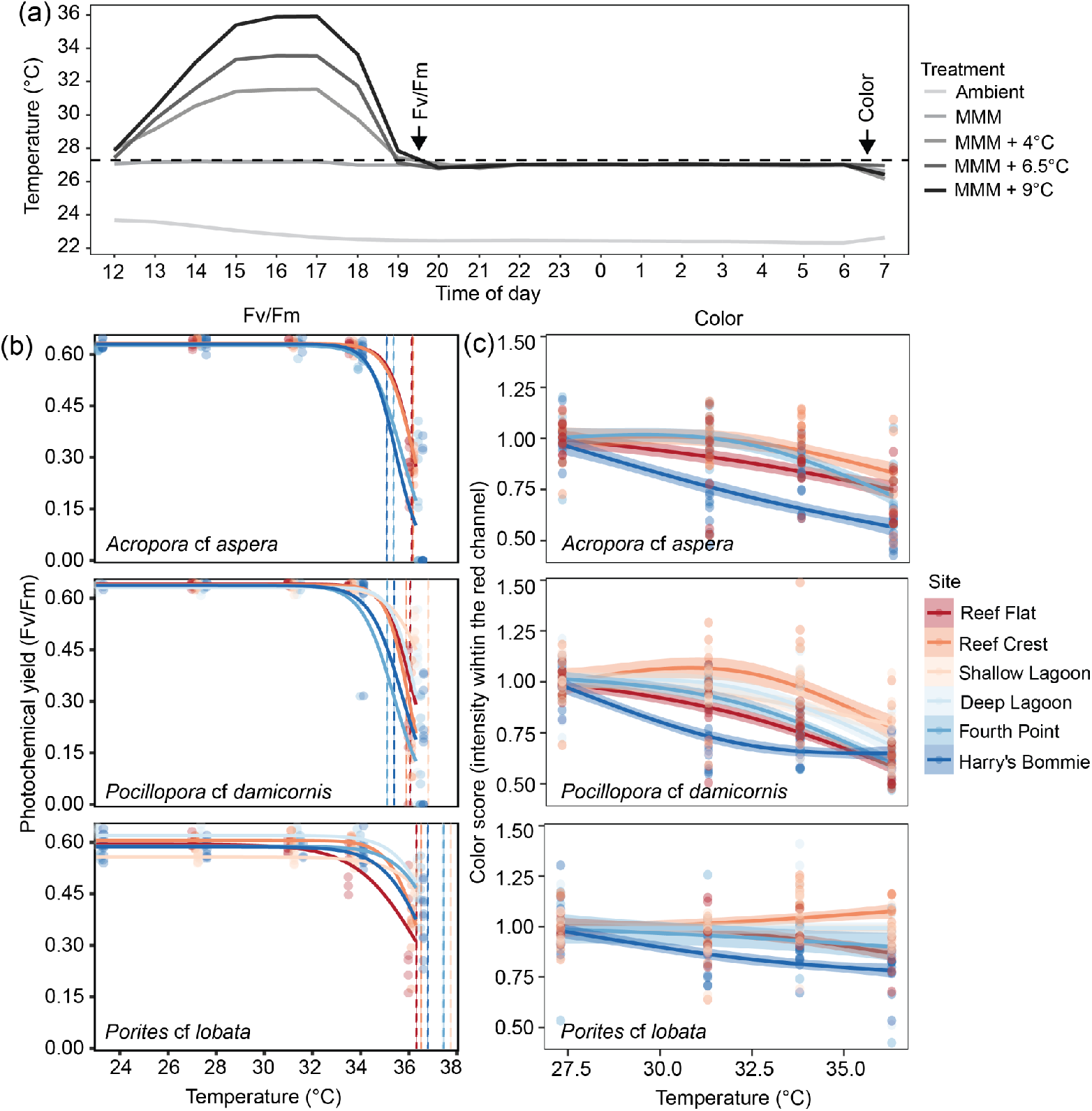
Physiological performance and coral heat tolerance across sites and species. (a) Hourly temperature measurements of the standardized experimental heat stress assays, with arrows indicating when physiological measurements were determined. MMM (dashed horizontal line) is the region’s climatological maximum monthly mean (27.3°C). (b) Three-parameter log-logistic dose response curves were fitted for F_v_/F_m_ measurements in response to temperature, where points indicate individual measures for coral genets (n= 5–10) in each treatment. Dashed vertical lines indicate the effective dose 50 (ED50). (c) Generalized additive mixed models were fitted for color score in response to temperature, where points indicate individual measures for coral genets in each treatment and confidence bands indicate 95% confidence intervals.

Photochemical yield (F_v_/F_m_) was significantly influenced by the three-way interaction of treatment, genus and site (X^2^ = 87.7, p < 0.0001). Pairwise comparisons generally revealed significant declines in F_v_/F_m_ in the hottest treatment (MMM+9°C) across all genera (Figure 4b; Figure S2). Across all sites, we measured a 1.45°C range in ED50 between the least and most heat tolerant genera (*Acropora* < *Pocillopora* < *Porites*) (Figure S2). *Acropora* was the least heat tolerant, with an ED50 of 35.65°C (95%CI: 35.2–36.1), followed by *Pocillopora* at 35.81°C (95%CI: 35.5–36.1) (Figure S2). The most heat tolerant genus was *Porites*, with an ED50 of 37.1°C (95%CI: 36.4–37.8) (Figure S2).

For each site, post hoc analyses revealed interesting patterns in baseline (i.e., ambient) photophysiology across the genera. At both reef slope sites (Harry’s Bommie and Fourth Point) and the Shallow Lagoon, photochemical yield was greater in *Pocillopora* compared to *Porites* (p < 0.02, p < 0.008, and p < 0.02, respectively), whereas at the Deep Lagoon, Reef Crest and Reef Flat, photochemical yield did not differ between the three species. *Acropora* did not differ in photochemical yield between sites or species at ambient (p > 0.98).

Significant differences in ED50 within each genus were uncovered across sites (Figure 4b). For *Acropora*, ED50 (mean; 95%CI) differed by 1.10°C between the least and most heat tolerant sites (HB < FP < RF < RC), ranging from 35.1°C (95%CI: 34.4–35.8) and 35.4°C (95%CI: 34.9– 35.8) at the less variable (i.e., <1.34°C day^-1^) reef slope sites Harry’s Bommie and Fourth Point, respectively. ED50’s increased up to 36.1°C (95%CI: 35.6–36.7) at the Reef Flat and peaked at 36.2°C (95%CI: 35.7–36.6) at the Reef Crest (Figures 4–5). *Pocillopora* exhibited the greatest range in ED50 (1.71°C) across the sites (FP < HB < RC < RF < DL < SL) (Figures 4–5). Again, *Pocillopora* from Fourth Point (ED50: 35.1°C; 95%CI: 34.6–35.6) and Harry’s Bommie (ED50: 35.4°C; 95%CI: 34.8–35.9) had the lowest thermal tolerance (Figures 4–5). Interestingly, *Pocillopora* from the sites with the greatest diel thermal variability (Reef Flat and Reef Crest) were not the most thermally tolerant (ED50: 35.9–36.1°C) (Figures 4–5). Instead, *Pocillopora* from the site with intermediate diel temperature variability (Shallow Lagoon) had the greatest thermal tolerance. (ED50:36.81°C; 95%CI:34.9–38.6) (Figures 4–5). For *Porites*, the ED50 ranged 1.43°C between the least and most heat tolerant sites (RF < RC < HB < DL < FP < SL). Surprisingly, the less variable sites Fourth Point (ED50: 37.45°C; 95%CI: 33.6–41.3) and Deep Lagoon (ED50: 37.40°C; 95%CI: 34.8–40.0) were associated with greater thermal tolerance for *Porites* than the two most variable sites, Reef Flat (ED50:36.3°C; 95%CI: 35.7–36.9) and Reef Crest (ED50:36.5°C; 95%CI: 35.8–37.2) (Figures 4–5). Yet, heat tolerance in *Porites* (ED50: 37.75°C; 95%CI: 30.8–44.7) was again greatest at the site with intermediate diel temperature variability (Shallow Lagoon) (Figures 4–5).

### Bleaching severity varied by genus and site under acute heat stress

Significant differences in coral color between genera and sites were found at MMM (X^2^ = 20.1, p = 0.009) (Figure S2). Pairwise comparisons revealed no differences in *Acropora* pigmentation across sites (p > 0.79) (Figure S2). Yet, *Pocillopora* originating from the Reef Crest and Deep Lagoon were less pigmented than corals from all other sites (p < 0.04), and *Porites* originating from the most (Reef Flat) and least thermally variable habitats (Harry’s Bommie) were significantly more pigmented than *Porites* originating from intermediate sites (Shallow Lagoon and Deep Lagoon) (p < 0.02) (Figure S2).

Normalized coral color was significantly influenced by the three-way interaction of treatment, genus and site (X^2^ = 35.7, p = 0.058) (Figure 4c). The severity of the bleaching response differed between genera. For *Acropora*, corals from the least (Harry’s Bommie) and most (Reef Flat) thermally variable sites showed the sharpest declines in pigmentation (Figure 4c). Pairwise comparisons revealed significant declines in color between MMM and all other temperatures in corals from Harry’s Bommie (p < 0.0001), whereas for the Reef Flat corals, there was no difference between MMM and MMM+4°C (p = 0.22), yet significant declines at MMM+6.5°C and MMM+9°C (p < 0.04) (Figure 4c). *Acropora* from the Reef Crest and Fourth Point did not show a decline in color between MMM, MMM+4°C, or MMM+6.5°C (p > 0.42), however, there was a significant decline in color at MMM+9°C in corals from both sites (p < 0.0001) (Figure 4c).

For *Pocillopora*, corals from Harry’s Bommie showed the sharpest decline in pigmentation, with pairwise comparisons revealing significant initial losses in color between MMM and MMM+4°C (p < 0.0001), yet no further declines between MMM+4°C, MMM+6.5°C, and MMM+9°C (p > 0.42). *Pocillopora* from the Reef Flat and Fourth Point showed similar trends (Figure 4c). Corals from the Reef Flat showed significant initial losses in pigmentation between MMM and MMM+4°C (p= 0.02) as well as further declines between MMM+6.5°C and MMM+9°C (p < 0.0001), whereas for corals from Fourth Point, there were no initial losses in color between

MMM and MMM+4°C (p = 0.99), yet stepwise declines between all other temperature comparisons (p < 0.0001) (Figure 4c). For *Pocillopora* from the Deep Lagoon, Reef Crest and Shallow Lagoon, there were no differences in color between MMM and MMM+4°C or MMM+6.5°C (p > 0.13), yet significant declines at MMM+9°C (p < 0.0001) (Figure 4c).

Remarkably, *Porites* exhibited no significant losses in pigmentation across temperatures at any site (p > 0.05) except Harry’s Bommie, where bleaching severity significantly increased between MMM and all other temperatures (p < 0.02) (Figure 4c). Interestingly, for *Porites* from the Reef Crest, pigmentation was lowest in the coolest treatment (MMM), significantly increasing at MMM+6.5°C (p < 0.0001) (Figure 4c).

### Diel temperature variability was the best predictor of coral heat tolerance

Variation in ED50 was explored against *in situ* temperature conditions recorded September 2015–August 2016. A non-linear model that included the individual effect of mean diel temperature amplitude resulted in the best prediction, as determined by the lowest AICc, explaining 28% of the variation. Thermal tolerance increased from a mean diel temperature amplitude of 1.06°C day^-1^, reaching the vertex or optimum at 2.2°C day^-1^, and decreasing when amplitude exceeded 2.81°C day^-1^ (Figure 5a). While the inclusion of coral species did improve explanatory power by ∼10%, it did not result in the best model (i.e., lowest AICc). Further, other environmental predictors and interactions including mean, minimum, and/or maximum temperature as well as maximum DHW experienced in 2016 or 2020 were explored and none improved the model.

**Figure 5.**
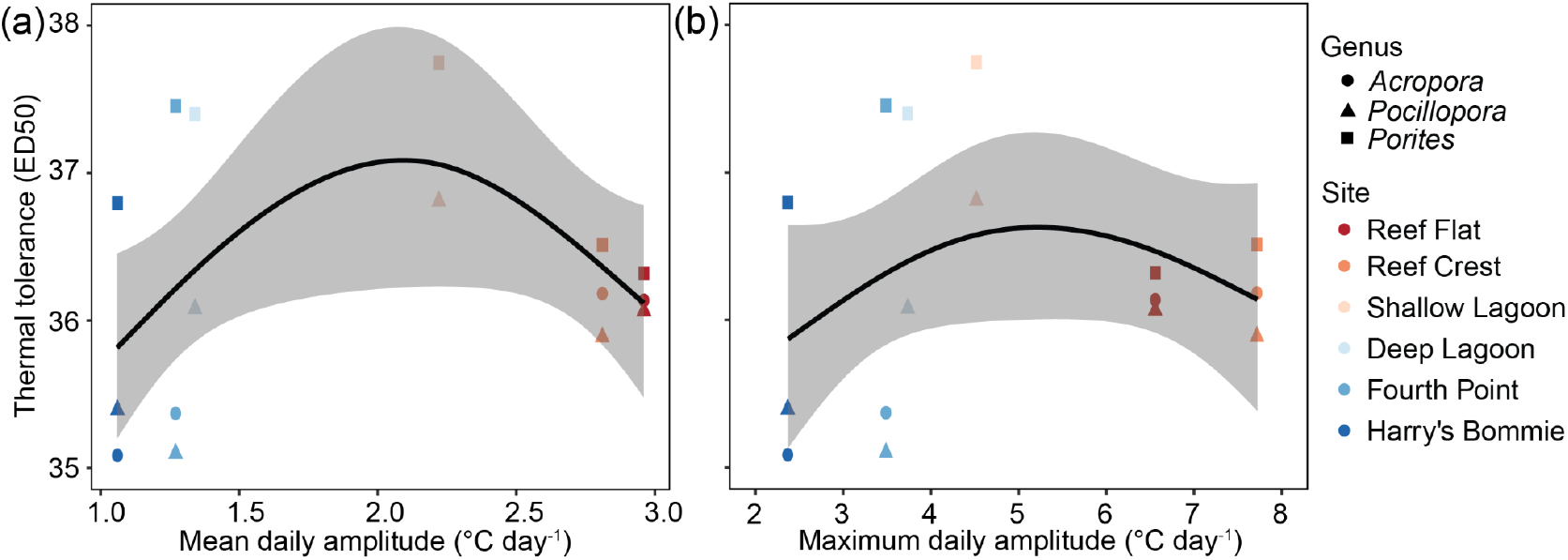
Relationship between coral thermal tolerance and thermal variability. (a) Relationship between coral thermotolerance (i.e., F_v_/F_m_ ED50) and mean diel temperature variability for the best generalized additive model (GAM). (b) Relationship between coral thermotolerance and maximum diel temperature variability. Points indicate ED50 for each species and each site and confidence bands indicate 95% confidence intervals.

To increase comparability with other studies investigating diel temperature variability on coral thermal tolerance, we also plotted variation in ED50 against maximum diel temperature amplitude. A non-linear model that included the individual effect of maximum diel temperature amplitude resulted in the second best prediction after mean diel amplitude, explaining 17.5% of the variation (Figure 5b). The trend mirrored that for mean diel temperature variability, with a maxima observed at intermediate variability. The inclusion of coral species did not improve explanatory power.

## Discussion

### Relative heat tolerance is driven by the amplitude of daily temperature variability

By investigating the thermotolerance of three species of corals originating from six reef habitats representing a wide range of diel temperature variability, we revealed a consistent parabolic response to increasing thermal variability across species, whereby coral thermotolerance was highest on reefs with intermediate temperature variability. Critically, the consistent response across corals of diverse life-history strategies (competitive, weedy and stress-tolerant) and phylogenies (Complexa and Robusta) is powerful, as it expands upon previous studies investigating one (Kenkel & Matz, 2016; Palumbi et al., 2014; Voolstra et al., 2020) or two species (Klepac & Barshis, 2020; Schoepf et al., 2015) in isolation. Further, the investigation of thermotolerance across a rigorously quantified thermal spectrum demonstrates that a larger range in variability (i.e., beyond two habitats) may be needed to reveal the full influence of variability on thermal tolerance. For example, focusing on only the least and most variable habitats at Heron Reef would have led to an erroneous conclusion that increasing temperature variability decreases heat tolerance for *P*. cf. *lobata*, as this species was 0.5°C less thermotolerant at the most variable habitat. Yet at intermediate variability, *P*. cf. *lobata* were 1°C more thermotolerant than the least variable habitat, indicating that there is an optimal intermediate priming exposure for this species as well, beyond which becomes too physiologically stressful for corals within the most extreme environments. Elevated thermal tolerance in corals exposed to intermediate diel temperature variability also occurs in *Porites lobata* from the pools of Ofu, American Sāmoa, where corals native to the moderately variable pool have higher thermotolerance than corals from the low or highly variable pools (Klepac & Barshis, 2022). However, these findings are in contrast to a number of earlier studies on *Acropora hyacinthus* from these same American Sāmoan pools, which consistently exhibit greater heat tolerance in the highly variable pools (Palumbi et al., 2014; Thomas et al., 2018). These discrepancies highlight the need to evaluate thermal tolerance using standardized comparisons across diverse species (Grottoli et al., 2021) and indeed, the examination of three diverse coral taxa in this study demonstrates a consistent response to temperature variability across species assemblages. Ultimately, we build support for observations that not all temperature regimes result in greater thermotolerance (Klepac & Barshis, 2022; Schoepf et al., 2019), and provide new evidence that this mechanism is congruent across diverse coral species.

### Elevated thermal tolerance does not protect against bleaching and mortality during marine heatwaves

Temperature regimes that expose corals to sub-lethal heat stress have been recognized as a mechanism to increase the physiological preparation for marine heatwaves (Ainsworth et al., 2016; Safaie et al., 2018; Sully et al., 2019; Thomas et al., 2018), and coral populations acclimated and/or adapted to variable thermal conditions are posited to be a source of climate resilience (e.g., (Palumbi et al., 2014)). However, during the 2020 marine heatwave that hit Heron Reef, coral bleaching and mortality were highest in the most thermally variable sites relative to the least thermally variable sites (Ainsworth et al., 2021; Brown et al., 2022). For example, branching *Acropora* exhibited 50-fold lower symbiont densities at the most thermally variable site relative to those from the least variable habitat (Ainsworth et al., 2021), resulting in a loss of nearly all branching *Acropora* (Brown et al., 2022). This was despite *Acropora* native to these same sites exhibiting 1.1°C higher bleaching thresholds than conspecifics from the least variable habitats, suggesting that even a 1°C advantage in thermotolerance gained from lifelong exposure to thermally variable conditions does not protect against current marine heatwaves. On the other hand, as our study was conducted 2 years after a marine heatwave, significant coral mortality stemming from this event may have resulted in a strong selection for the most thermally-tolerant genotypes (Burgess et al., 2021; Marzonie et al., 2022; Sampayo & Ridgway, 2008). Additional studies are needed to determine the mechanisms driving elevated heat tolerance in variable and moderately variable environments, such as genetic variability in the host and/or associated Symbiodiniaceae (e.g., (Oliver & Palumbi, 2011b)), and whether genotypes capable of surviving extreme temperature variability can also withstand prolonged and repeated marine heatwaves. Nonetheless, our results add to a body of evidence that corals exposed to extreme thermal regimes are unable to cope with the additional heat stress of marine heatwaves superimposed on top of thermally variable conditions (Ainsworth et al., 2021; Brown et al., 2022; Schoepf et al., 2015, 2020).

As climate change intensifies, the response of corals and trajectories of ecosystems are becoming more contingent on previous marine heatwaves (e.g., (Hughes et al., 2019)). There is growing evidence that surviving corals can acclimatize (i.e., acquire stress tolerance through hardening) or sensitize (i.e., accumulate stress leading to weakening) via the environmental memory of thermal stress (Brown & Barott, 2022; Hackerott et al., 2021). Whether the corals investigated in this study are acclimatizing or suffering from long-term damage from the 2020 marine heatwave, and if this environmental memory influenced thermotolerance, cannot be determined and is likely species-specific (Evensen et al., 2022; Marzonie et al., 2022). However, it is plausible that the lower thermotolerance of *P*. cf. *lobata* from the most thermally variable habitat, which experienced disproportionately higher heat stress during the 2020 marine heatwave (Brown et al., 2022), is a result of stress accumulation across repetitive marine heatwaves — similar to the recent findings of reduced thermal tolerance of *Porites* from the Red Sea and American Sāmoa following heatwaves (Evensen et al., 2022; Klepac & Barshis, 2020, 2022). Conversely, greater thermotolerance of *P*. cf. *lobata* from the least thermally variable habitats could have stemmed from a magnitude and duration of heat stress during the 2020 heatwave that promoted stress hardening of this species (Hackerott et al., 2021; Marzonie et al., 2022). Ultimately, it remains inconclusive whether the patterns observed here are related to acclimatization or sensitization, but a better understanding of these processes is key to predicting the future of coral reefs in warming oceans and are an important avenue of future studies (Brown & Barott, 2022; Hackerott et al., 2021).

### Local habitat heterogeneity rivals regional differences in coral thermal tolerance

The range in coral thermotolerance across geomorphological zones within the single reef system of Heron Reef (<5 km) were as high as differences observed across vast latitudinal gradients in the Caribbean (∼300km, Florida Reef Tract) (Cunning et al., 2021), eastern Australia (∼860 km, Coral Sea) (Marzonie et al., 2022), and the Red Sea (∼900 km) (Evensen et al., 2022). For example, *A*. cf. *humilis, P. verrucosa* and *P. meandrina* on reefs across the Coral Sea — spanning 7.7 degrees of latitude and corresponding with a 1.6°C gradient in MMM — led to 0.85°C to 1.89°C range in heat tolerance across sites (Marzonie et al., 2022). Similarly, *A. hemprichii, P. verrucosa*, and *P. lobata* from reefs across the Red Sea spanning 17 degrees of latitude and a 3.7°C gradient in MMM displayed a 1.1°C to 1.6°C range in thermotolerance (Evensen et al., 2022). In this study, we observed a comparable range of thermotolerance (1.1– 1.71°C) across the six reef habitats investigated, suggesting fine-scale temperature heterogeneity can increase thermal thresholds similar to large-scale differences in temperature across latitudinal gradients. While microhabitats are known to shape patterns in coral thermotolerance across small spatial scales (Schoepf et al., 2015; Thomas et al., 2018; Voolstra et al., 2020), our standardized comparison across three corals with distinct life-history strategies allows for the comparison of these traits for the first time across regions and spatial scales. For example, we observed a greater range in thermal tolerance in *P*. cf. *damicornis* (1.71°C) across Heron Reef (<5 km) than congeners *P. meandrina* (1.15°C) or *P. verrucosa* (0.85°C) across the Coral Sea (∼860 km) and *P. verrucosa* (1.55°C) across the Red Sea (∼900 km), whereas for acroporids, there was a slightly lower range in thermal tolerance across Heron Reef (0.9°C) than the Red Sea (1.1°C) but half the range in the Coral Sea (1.89°C). These differential patterns in thermal tolerance are likely a result of: (i) adaptation to latitudinal thermal regimes over evolutionary time (Dixon et al., 2015; Osman et al., 2018), (ii) variation in host or symbiont communities (Burgess et al., 2021; Oliver & Palumbi, 2011b; Sampayo & Ridgway, 2008) and/or (iii) recent exposure to marine heatwaves (Evensen et al., 2022; Hughes et al., 2019; Marzonie et al., 2022). This is encouraging, particularly as gene flow from the thermally variable habitats could increase thermal tolerance within the least thermally variable habitats within the same reef system. Yet, investigations into *P. damicornis* across the thermally variable reef flat and thermally stable reef slope of Heron Island have indicated that there is limited gene flow between these populations, which may be a result of differences in genetic makeup and/or maladaptation to environmental conditions, given that the small spatial distances between reef habitats (<100 m) would not be expected to restrict dispersal (van Oppen et al., 2018). Given limited gene flow, human interventions like assisted gene flow and/or selective breeding may be a viable strategy to increase heat tolerance of certain coral populations across thermally-distinct reef habitats (Van Oppen et al., 2017), offering an easier and safer alternative to moving thermotolerant corals across latitudes (e.g., (Dixon et al., 2015)). Indeed, *P. damicornis* can survive transplantation from the most to least thermally variable habitats at Heron Reef, and even retains greater heat tolerance than native conspecifics for at least 18 months following transplantation (Marhoefer et al., 2021). While it may be impossible to perform these tasks across the entirety of the Great Barrier Reef, reefs such as Heron Reef, which is a high-value tourism and world-class research destination, are ideal candidates for such stewardship to increase coral resilience in a changing climate.

## Conclusions

The results of this study highlight that fine-scale temperature heterogeneity can increase coral thermal thresholds in diverse coral lineages, and reveal that there is an optimal priming exposure at intermediate temperature variability that leads to maximal thermotolerance. Greater thermal tolerance, however, does not necessarily translate into greater community resilience during marine heatwaves, as the coral communities that had higher bleaching thresholds experienced more prevalent and severe bleaching and greater declines in hard coral cover following the 2020 heatwave (Ainsworth et al., 2021; Brown et al., 2022). This suggests that elevated heat tolerance gained from life-long exposure to sub-lethal thermal variability already appears ineffective against current levels of ocean warming. Encouragingly, the range in coral thermotolerance across geomorphological zones within a single reef system (< 5 km) were as large as differences observed across vast latitudinal gradients (>300 km), and future studies could investigate the mechanisms (e.g., physiological plasticity, constitutive upregulation of stress-response genes, and/or epigenetic modifications) enabling these corals to develop resistance to acute heat stress. Further, while co-occurring environmental conditions (e.g., pCO_2_, oxygen, and irradiance) were not quantified in this study, pCO_2_ fluctuations and irradiance are known to differ across these same habitats (Brown & Mello-Athayde, 2022), and the combined effect of ocean warming and deoxygenation can lower the thermal threshold of some corals (Alderdice et al., 2022). As such, more research is needed to understand the interactions between physicochemical conditions that co-occur within thermally variable habitats and their influence on coral thermal tolerance. To encourage the best future for coral reefs, the potential for assisted gene flow to increase heat tolerance of coral populations should continue to be explored, while concurrently adopting strict global policies to limit climate-induced temperature increases to 1.5°C (Hoegh-Guldberg et al., 2019).

