## Supplementary materials for "Maximal coral thermal tolerance is found at intermediate diel temperature variability"

### Methods

#### Image analysis of coral color, a proxy for bleaching severity

Coral color was determined from each photograph in a semi-automated manner. Each photograph was first cropped to a standard size to remove excess background via a custom automated batch script in Adobe Photoshop (Version 21.1.2). Photographs were then loaded into ImageJ (v1.53c (Schneider et al., 2012)), and the performance of 16 built-in segmentation models were tested on a sub-sample of coral images to remove the background of the cropped image, which was turned to black, thus leaving only the coral fragment. The segmentation model that best segmented all coral fragments from the background effectively and with limited coral fragment cut off (Model Li) was then implemented on all images, which were batch processed using a custom image segmentation macro script modified from (Strock, 2021). Once segmented, the script then extracted red pixel intensity of the fragment in RGB, HSB, and LAB color spaces. Finally, the mean red pixel intensity of the red color standard from the original (unsegmented) images was extracted from a region of interest drawn by hand in ImageJ. Pixel intensities of the coral and corresponding red standard were then converted to a ‘darkness’ score by subtracting the red channel 'brightness’ from the maximum value (255). The mean red channel darkness of each coral was then normalized by dividing by the mean red pixel darkness of the red color standard from the same photograph. These normalized color values were then used to calculate the changes in bleaching severity between species, sites and treatments. For visualization, the normalized color values were divided by the mean color under ambient (MMM) conditions for that species.

#### Statistical analyses

We fit all possible model combinations using the gam function from the package *mgcv* (Wood, 2006). The model structure was developed using a stepwise procedure, where models were compared and selected using the Akaike information criterion for small sample sizes (AICc) and the model with the lowest AICc was selected as the best model (Dove et al., 2020). Smooth terms were fit using thin plate regression splines (tp) and the number of knots were restricted (k =3) to avoid overfitting.

**Supplementary figures and tables**

**Table S1.** Details of coral collections and standardized short term heat stress (STHS) assays. Two treatments (n = 1 tank) were performed per day and each site was done in isolation, with treatments tested each day selected randomly. Treatment temperatures indicated are relative to the climatological maximum monthly mean (MMM) of 27.3°C.

| Site | Collection date | STHS date | Time STHS start | Time of photochemical yield ($F_v/F_m$) | Time of photograph for color score |
|---|---|---|---|---|---|
| Harry's Bommie (HB) | October 6, 2022 | Oct 7–Oct 8: +6.5°C & +9°C | 12:00 | 19:00 | 07:00 |
| | | Oct 8–Oct 9: MMM & +4°C | 12:00 | 19:00 | 07:00 |
| Fourth Point (FP) | October 4, 2022 | Oct 5–Oct 6: MMM & +4°C | 12:00 | 19:00 | 07:03 |
| | | Oct 6–Oct 7: +6.5°C & +9°C | 12:20 | 19:20 | 07:05 |
| Deep Lagoon (DL) | September 28, 2022 | Sept 29–30: MMM & +4°C | 12:00 | 19:00 | 06:45 |
| | | Sept 30–Oct 1: +6.5°C & +9°C | 12:00 | 19:00 | 07:15 |
| Shallow Lagoon (SL) | September 27, 2022 | Sept 27–28: +4.5°C & +9°C | 13:10 | 20:10 | 07:00 |
| | | Sept 28–29: MMM & +6.5°C | 12:15 | 19:15 | 07:00 |
| Reef Crest (RC) | October 2, 2022 | Oct 3–Oct 4: MMM & +4°C | 12:00 | 18:57 | 07:00 |
| | | Oct 4–Oct 5: +6.5°C & +9°C | 12:20 | 19:20 | 07:00 |
| Reef Flat (RF) | September 30, 2022 | Oct 1–2: +6.5°C & +9°C | 12:00 | 19:00 | 07:00 |
| | | Oct 2–3: MMM & +4°C | 12:00 | 19:00 | 07:00 |

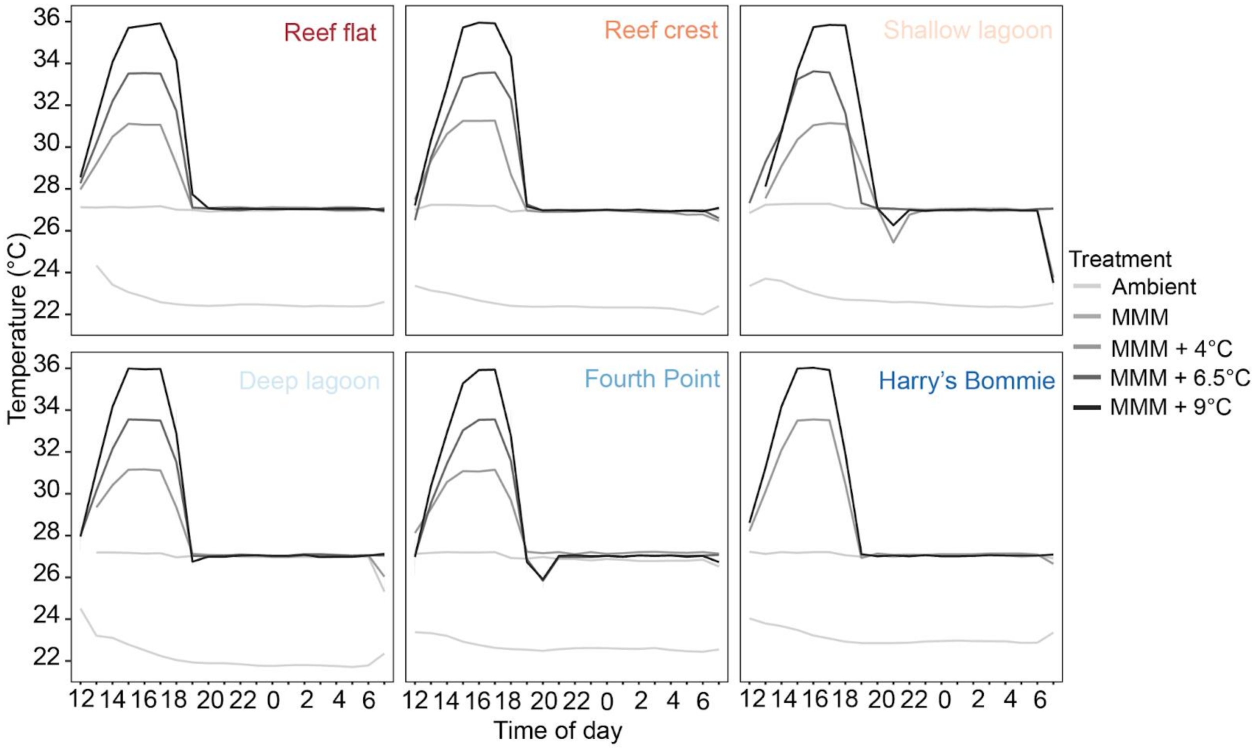


**Figure S1.** Hourly temperature profiles across experimental assays.

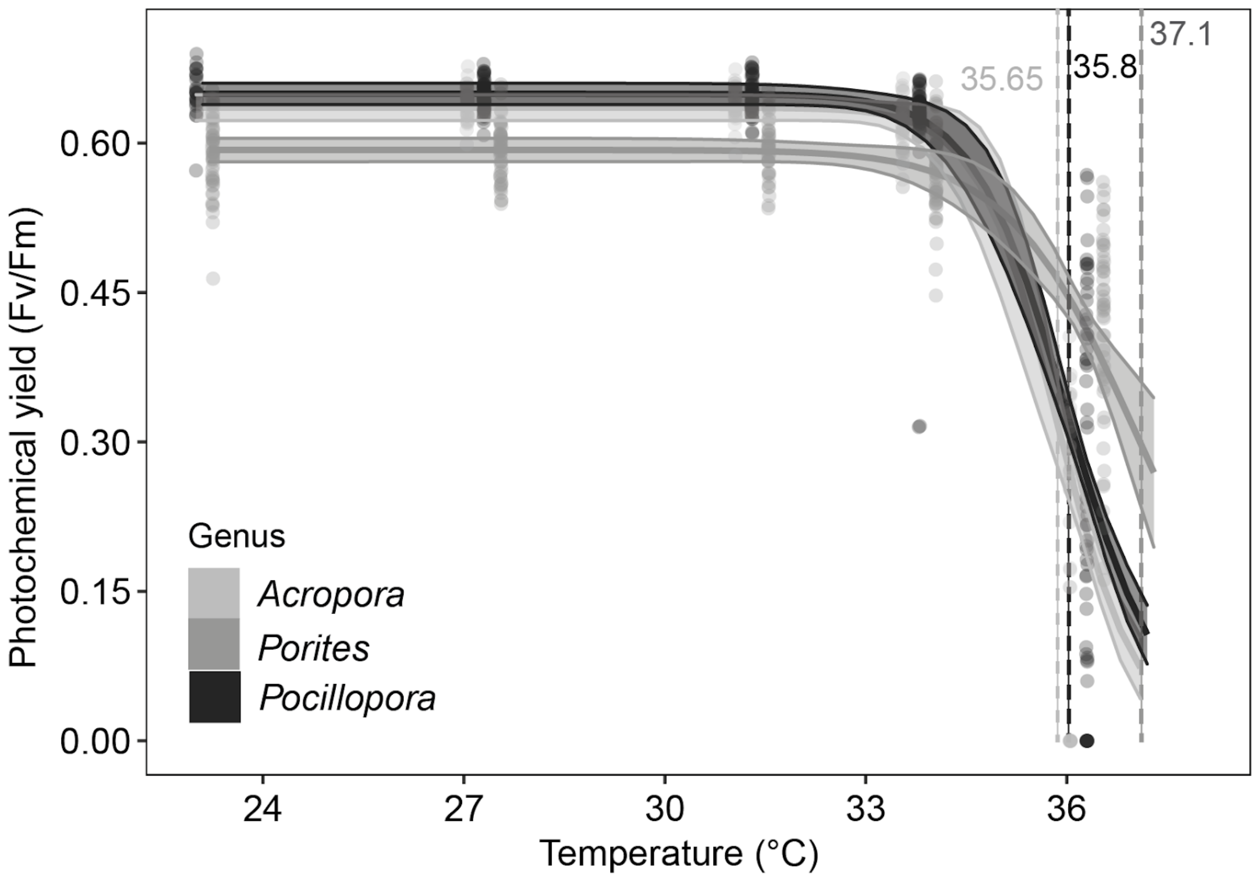


**Figure S2.** Coral photophysiological performance across genera. Points indicate measures of $F_v/F_m$ for individual coral genets (n= 5–10) in each treatment. Confidence bands indicate 95% confidence intervals. Dashed vertical lines indicate the effective dose 50 (ED50).

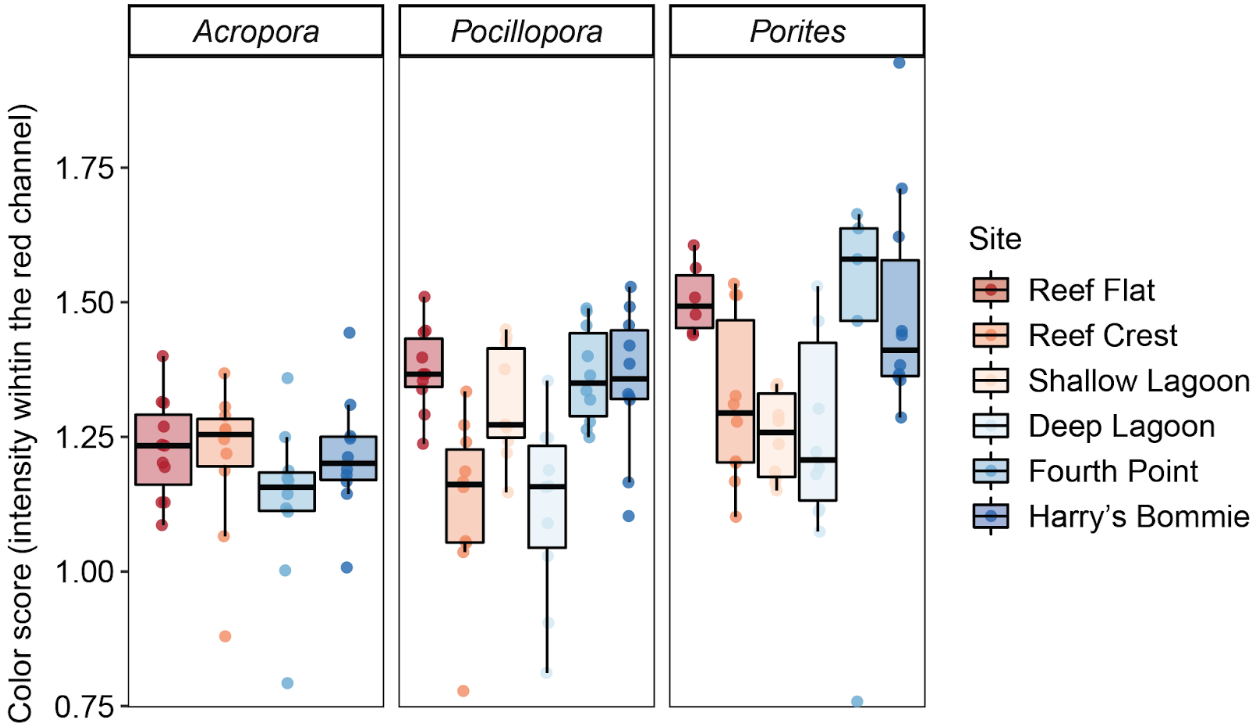


**Figure S3.** Differences in coral color between genera and sites at the control temperature (MMM; 27.3°C). Color score was determined from the red channel pixel intensities of normalized RGB photographs. Boxplots display the minimum, 25$^{th}$ percentile, median, 75$^{th}$ percentile, and maximum, where points indicate measures for individual coral genets (n= 5–10) at each site.
